# The genetic interaction of *REVOLUTA* and *WRKY53* links plant development, senescence, and immune responses

**DOI:** 10.1101/2021.07.27.454007

**Authors:** Justine Bresson, Jasmin Doll, François Vasseur, Mark Stahl, Edda von Roepenack-Lahaye, Joachim Kilian, Bettina Stadelhofer, James M. Kremer, Dagmar Kolb, Stephan Wenkel, Ulrike Zentgraf

## Abstract

In annual plants, tight coordination of successive developmental events is of primary importance to optimize performance under fluctuating environmental conditions. The recent finding of the genetic interaction of *WRKY53*, a key senescence-related gene with *REVOLUTA*, a master regulator of early leaf patterning, raises the question of how early and late developmental events are connected. Here, we investigated the developmental and metabolic consequences of an alteration of the *REVOLUTA* and *WRKY53* gene expression, from seedling to fruiting. Our results show that *REVOLUTA* critically controls late developmental phases and reproduction while inversely *WRKY53* determines vegetative growth at early developmental stages. We further show that these regulators of distinct developmental phases frequently, but not continuously, interact throughout ontogeny and demonstrated that their genetic interaction is mediated by the salicylic acid (SA). Moreover, we showed that *REVOLUTA* and *WRKY53* are keys regulatory nodes of development and plant immunity thought their role in SA metabolic pathways, which also highlights the role of *REV* in pathogen defence. Together, our findings demonstrate how late and early developmental events are tightly intertwined by molecular hubs. These hubs interact with each other throughout ontogeny, and participate to the interplay between plant development and immunity.

## Introduction

The life cycle of flowering plants can be considered as a series of distinct growth phases driven by developmental genetic programs that integrate both environmental and endogenous stimuli. In annual plants, leaf senescence, defined as the last developmental stage, is often considered as an essential trait of plant adaptation to its biotic and abiotic environment [1]. Leaf senescence is notably of utmost importance in the crosstalk between developmental, abiotic stress and immune responses, and influences plant productivity and fitness, as well as resistance to pathogens [2]. However, how senescence-related genes connect plant development, abiotic stress and immune responses still remains unclear. Moreover, this raises questions how early and late developmental events are coordinated by molecular hubs expressed throughout the life cycle.

The onset, progression and completion of leaf senescence are tightly regulated and depend on both plant age and growth environment. The senescence process occurs in an orderly manner, and without exogenous stress it mainly depends on the integration of age information at leaf and whole-plant levels [3]. In *Arabidopsis thaliana*, senescence is initiated as soon as full expansion of the leaves is reached and usually coincides with the transition from vegetative to reproductive growth [4]. During senescence progression, sequential changes arise coordinately in plant physiology and metabolism, and so implicate a large variety of genes involved for examples in the regulation of hormone, sugar and reactive oxygen species (ROS) levels [5]. The network of senescence-related genes initiates a well-orchestrated degradation of chlorophyll and other macromolecules, resulting in a sharp decrease of leaf photosynthetic activity [6]. When all essential nutrients have been remobilized to reach the reproductive parts of the plant, leaf death occurs as terminal phase of senescence [5,7].

Even though senescence is developmentally programmed, it can be strongly modulated by various exogenous factors. Stress, such as water stress or pathogen attack, can for example induce premature senescence as exit strategy to guarantee offspring under long-lasting unfavourable conditions [5]. Altogether, senescence is the result of a balance between developmental and environmental clues, integrating major transcriptional changes. In the case of pathogen infection, immune responses are induced and interfere with age-induced senescence signals, which can, in some cases, lead to a precocious senescence [2]. Interestingly, large-scale analyses of gene expression in senescing leaves of *A. thaliana* revealed that defence-related genes represent a significant portion of the leaf senescence transcriptome [8]. Indeed, it has been shown that their respective signalling pathways greatly overlap and several senescence-associated genes (SAGs) are activated during both development and defence [8,9]. A large fraction of the genes that operate at the nexus of development and defence encode proteins involved in hormonal signalling. For instance, jasmonic (JA) and salicylic (SA) acids have long been recognized as key hormones in the interconnection between age- and stress-induced senescence [10]. JA, SA as well abscisic acid (ABA) levels increase during age-related senescence [6] and signalling mutants are impaired in senescence onset and/or progression [11–13].

Transcription factors (TF) also play a pivotal role as they often act as regulatory nodes between signalling pathways and thus contribute to the fine-tuning of developmental and defence responses [2]. Among the most relevant TFs, the *WRKY* gene family constitutes the second largest group of the senescence transcriptome [14] and several *WRKYs* are also implicated in plant immunity [15]. Consistently, *WRKY53* acts at a convergence point between age-induced and stress-induced senescence [16,17]. For instance, *WRKY53* is known to be a positive regulator of senescence initiation and interacts with a large number of genes involved in senescence signalling, including other *WRKY* members and various SAGs [17,18]. Moreover, *WRKY53* has been shown to be an important component of defence signalling pathways in Arabidopsis [16]. Dual functionality in plant immunity was observed: while *wrky53* mutant plants had increased susceptibility toward *Pseudomonas syringae* [19,20], delayed symptoms were displayed during *Ralstonia solanacearum* infection [21].

Owing to its key multi-function, *WRKY53* expression is under a complex control. Its expression is notably modulated by the cellular redox state, SA and JA. Its activity can be modulated by phosphorylation by a mitogen-activated protein kinase kinase kinase (MEKK1) or by interaction with *EPITHIOSPECIFYING SENESCENCE REGULATOR* (*ESR/ESP*), which both have functions in pathogen responses [22,23]. Recently, *REVOLUTA* (*REV*), a member of the class III homeodomain leucine zipper (*HD-ZIPIII*) TF family, has been identified as a direct and positive regulator of *WRKY53* expression [24]. *REV* is known to have pleiotropic effects during plant development [25]. It mainly regulates polarity associated-growth processes during early leaf development but also controls the formation of floral meristems [26,27]. REV is part of a regulatory network of HD-ZIPIII factors, miRNAs and microProteins [27–29]. Interestingly, an additional role for *REV* in late leaf development has been recently reported since loss-of-function mutations in *REV* strongly delayed the onset of senescence, through the control of the expression of *WRKY53* and other SAGs like e.g. *MORE AXILLARY BRANCHES 2* (*MAX2*) by the REV protein [24,30]. This result pointed out the importance of *REV* in the genetic control of age-induced senescence. However, despite its key role in senescence initiation, the implication of the *REV-WRKY53* genetic interaction through plant life cycle and immunity remains unknown.

In this paper, we examined whether the genetic interaction between *REV* and *WRKY53* occurs from early to late development phases, and controls plant metabolism and immune responses. Combining ecophysiological, metabolic and molecular analyses, we examined the role of *REV* and *WRKY53* in the dynamics of *A. thaliana* growth and responses to pathogen attack. This study was conducted using four mutants affected in *REV* and/or *WRKY53* expression, which allow deciphering the specific role of each gene and their combined roles. Our results showed that although *WRKY53* is a regulator of late leaf development, it also determined vegetative growth evens at early developmental stages. Inversely, *REV* critically controlled late developmental phases and reproduction although it is known as a master regulator of early plant development and leaf patterning. Their genetic interaction fluctuated throughout ontogeny and dependent on environmental clues. Moreover, we showed that the *REVOLUTA*-*WRKY53* interaction is a key regulatory node of plant development and immunity through its role in SA metabolic pathways.

## Results

### *REV* and *WRKY53* control metabolic changes during senescence, especially related to SA metabolism

Senescence was dynamically analysed from flowering (first flower opens) until later developmental stages, ending with fruit ripening (first yellowing mature siliques) in wild-type plants (Col-0) as well as in four different mutants affected in *REV* and/or *WRKY53* expression: the mutants *rev5*, *wrky53* and *wrky53rev5* present single and double knock-out mutations [22,24], whereas *rev10-d* is a semi-dominant gain-of-function *REV* allele [27]. In all tested mutants, senescence was delayed after flowering, with a significant and strong deceleration 10 days after flowering, compared to Col-0 wild-type (Fig 1). In *rev5*, *wrky53* and *wrky53rev5* mutants, this delay was mainly due to postponed initiation of senescence in the first (oldest) leaves of the rosette (leaf position 1-8; S1 Fig). At later stages, the knock-out mutants reached similar percentages of senescent areas to Col-0, whereas *rev10-d* mutants were significantly less senescent until fruit ripening (*i.e*., mature silique stage; Fig 1). This strong delayed senescence in *rev10-d* was the result of a higher number of younger healthy leaves and not a delay in the onset of senescence (leaf position 16-20; S1 Fig).

**Fig 1.**
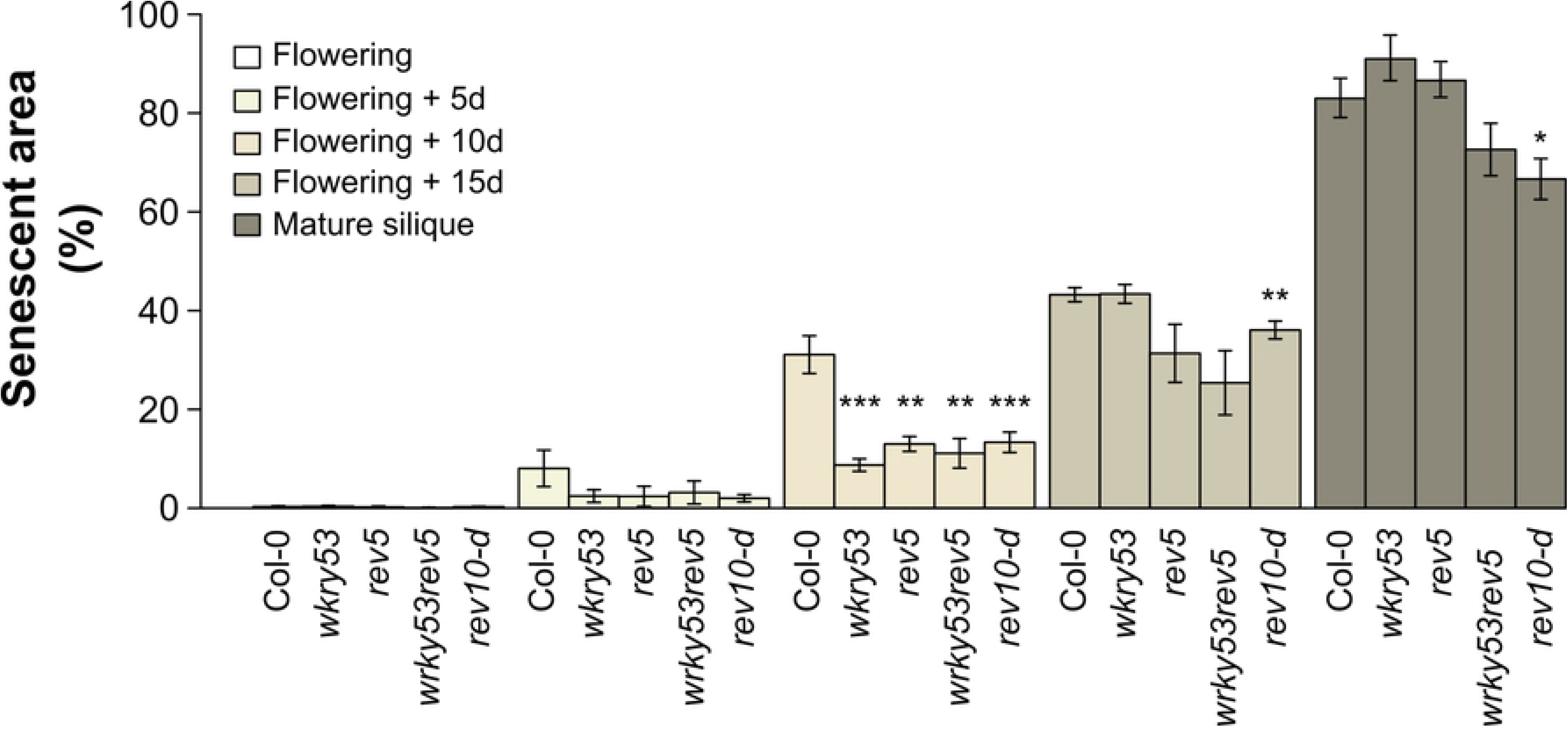
Analysis of senescence progression in the different *REV* and *WRKY53* mutants. Percentage of senescent area throughout the development from flowering to mature silique using chlorophyll fluorescence imaging. Senescent area was calculated by the ratio between pixel number for a photosynthetic efficiency *F*_v_/*F*_m_ ≤ 0.6 and the total pixel number of the rosette. Flowering was reached when the first flower was open. Mature silique stage was considered from the first yellowing siliques. Data are means (± SE) of 5-30 plants per genotype. Significant differences were analysed using Kruskal– Wallis tests: *: *P* ≤ 0.05, **: *P* ≤ 0.001, and ***: *P* ≤ 0.0001. See also S1 Fig.

To understand whether and how *REV* and *WRKY53* influence plant metabolism during senescence progression, we measured 22 metabolites from flowering until mature silique stage, including phytohormones, and related components, sugars, amino and organic acids. The heat map of response ratio between Col-0 and mutants, revealed that the hierarchical clustering of the different conditions (*i.e*., combining mutant and developmental stage) was not driven exclusively by plant development or by genotype but by a mixture between both (Fig 2A). At flowering, *rev5*, *wrky53* and *wrky53rev5* presented globally lower variations of response ratios (*i.e*. light colours) compared to *rev10-d* which presented a strong contrasting pattern, in which the majority of metabolites were found to be down-regulated, especially sugars and hormones (IAA, ABA, JA and SA-related components). The strongest observed dissimilarity in the response ratio of the mutants was found 15 days after flowering, while *rev5* at flowering was closely clustered to *rev10-d* at mature silique stage (Fig 2A). Throughout development, the double mutant *wrky53rev5* presented different patterns: it was closely clustered to *wrky53* at flowering and 5 days after flowering whereas it was more similar to *rev5* for the later stages.

**Fig 2.**
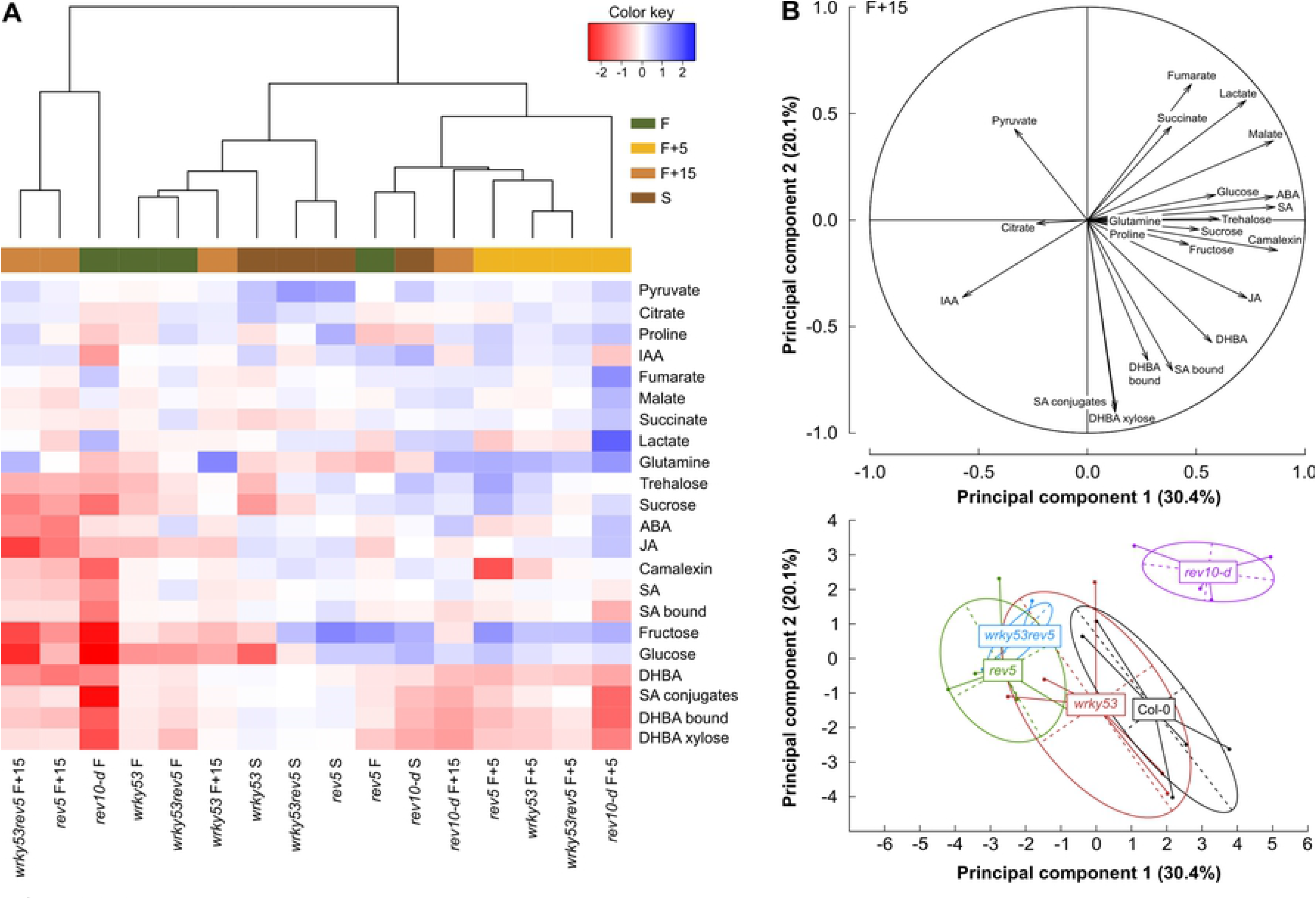
Metabolite patterns in *REV* and *WRKY53* mutants over development. **(A)** Heat map analysis of metabolites in *REV* and *WRKY53* mutants compared to wild-type Col-0. The row represents metabolites and the column displays the different genotype-stage combinations. The colour scale indicates the values of response ratio, calculated between the average of log-transformed values of Col-0 and a mutant at a given stage. Metabolite decreased is displayed in red, while metabolite increased is displayed in blue. White colour indicates no difference. The brightness of each colour corresponds to the magnitude of difference in the response ratio. The dendrogram represents the proximity between each genotype-stage metabolic pattern, as calculated from a hierarchical clustering analysis using Manhattan distance measures and Ward’s hierarchical clustering method. **(B)** Principal component analysis on metabolite abundance values at 15 days after flowering. **Top**: representation of metabolites on the two first principal components. **Bottom**: projection of individual plants with centres of gravity per genotype (n=5). See also Tables S1–S2 and S2 Fig. IAA = auxin, ABA = abscisic acid, JA = jasmonic acid, SA = salicylic acid, SA bound = total SA after hydrolysis - free SA, SA conjugates = SA-(C6)-glycosides, DHBA = dihydroxybenzoic acid; DHBA bound = Total DHBA after hydrolysis - free DHBA; F = flowering; d = days and S = mature silique stage.

The differences in the metabolic patterns of mutants compared to Col-0 at 15 days after flowering were analysed through a PCA of metabolites (Fig 2B; S1 Table). PC1, which accounted for 30.4 % of the total variance, was strongly explained by camalexin content, which is a phytoalexin involved in plant responses to pathogen infection and is linked to ROS and SA signalling [31]. PC1 was secondly explained by SA, ABA, malate and JA contents. Variations along PC1 were mainly driven by *rev5* and *wrky53rev5* genotypes (*P* < 0.01; ANOVA on PC coordinates), which exhibited lower contents of these metabolites compared to Col-0 (S2 A-E Figs), as illustrated by their respective distance from the centroid of Col-0 (Fig 2B, S1 Table). PC2, which accounted for 20.1 % of the total variance, was mainly explained by SA-related components, such as dihydroxybenzoic acid (DHBA)-xylose and DHBA-bound (total DHBA after hydrolyses - free DHBA), which are main SA catabolites in Arabidopsis [32] but also SA-conjugates (SA-(C6)-glycosides) and SA-bound (total SA after hydrolyses – free SA). Variations along PC2 were mainly driven by *rev10-d*, which presented the most significant genotypic effect (*P* < 0.01; ANOVA on PC coordinates; Fig 2B; S1 Table). Consistently, *rev10-d* displayed significant (*P* < 0.01) lower SA-conjugates and DHBA-xylose compared to Col-0 (S2 F-G Figs). Amongst all genotypes, *wrky53* displayed the least metabolic changes at 15 days after flowering compared to Col-0 (Fig 2B and S2 Fig). The analysis of the double mutant in senescing plants, by two-way ANOVA revealed an additive effect of *REV* and *WRKY53* mutations on SA and SA-related compound contents (S2 Table).

### The salicylic acid signalling pathway is interconnected to *REV* and *WRKY53* genes

In view of metabolic changes occurring in the mutants, we explored how *REV, WRKKY53*, and SA metabolism could be linked.

We first evaluated the modifications of the SA signalling pathway in the *REV* and *WRKY53* mutants through two key genes in SA biosynthesis and catabolism (Figs 3A-B). The *ISOCHORISMATE SYNTHASE 1* (*ICS1;* also known as *SALICYLIC ACID INDUCTION DEFICIENT 2*, *SID2*) gene, which is involved in SA biosynthetic process [33], presented an increasing expression during senescence progression in Col-0 (Figs 1 and 3A). This up-regulation was significantly reduced in *wrky53rev5* upon 5 days after flowering compared to Col-0, whereas *rev5* and *rev10-d* had a strong reduction at 15 days after flowering (Fig 3A). In contrast, *wrky53* did not show any significant difference in *ICS1* expression over time. The expression of the *SA 3-HYDROXYLASE* (*S3H*; also named *SAG108*) gene which is involved in SA catabolism [34], exhibited similar trends as per *ICS1*. The expression of the *S3H* gene in Col-0 was increased during senescence and expression in all mutants tended to be lower at 15 days after flowering, with a significant effect for *wrky53* and *rev5* (Fig 3B).

**Fig 3.**
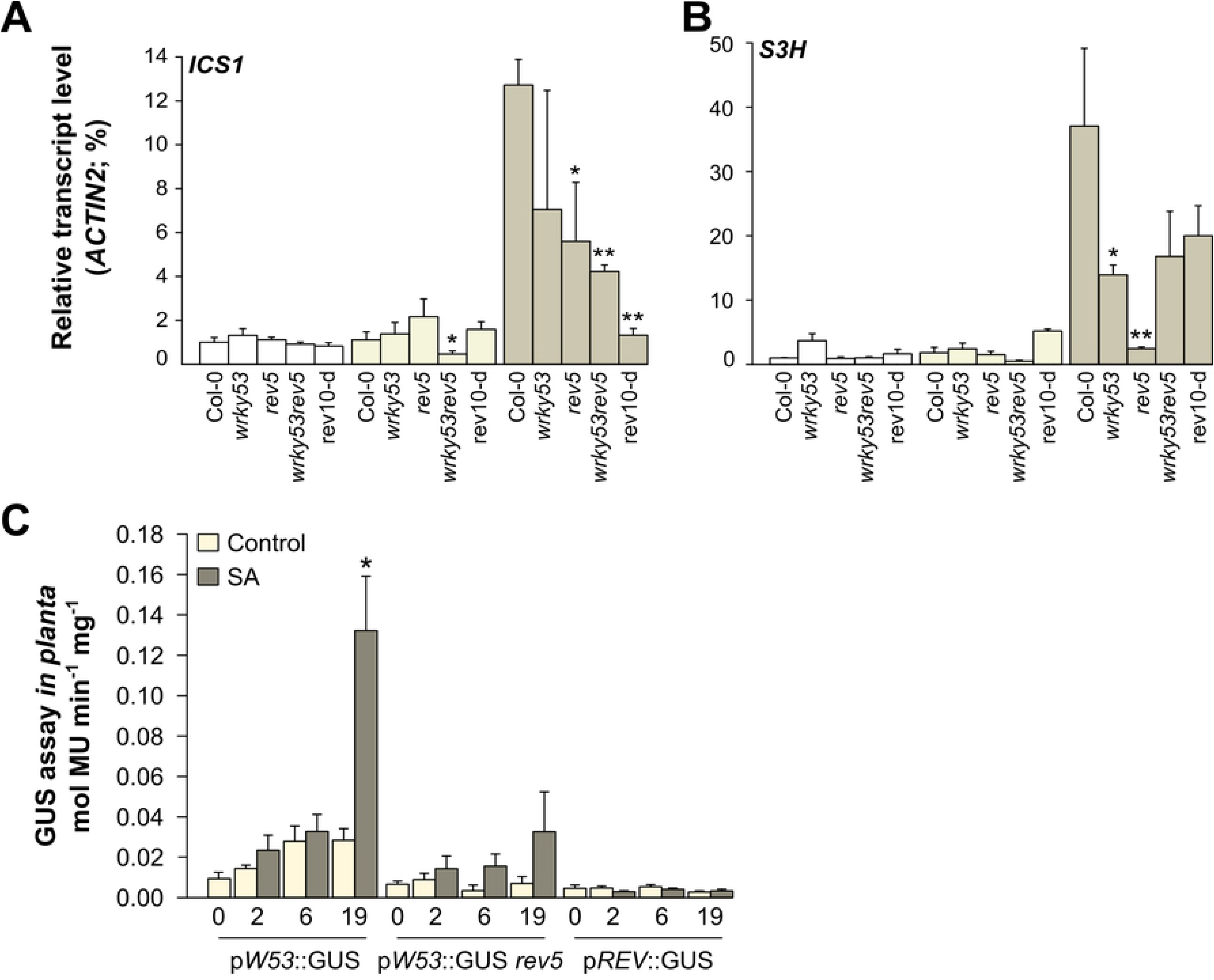
Genetic regulation of SA-related genes and promoter activities of *REV* and *WRKY53* after salicylic acid treatments. Gene expression analysis by qRT-PCR over development: (**A**) *ICS1* and (**B**) *S3H*. qRT-PCR was performed in leaf at position 10, pooled by 5 per genotype per stage. Transcriptional levels were calculated based on the comparative ΔΔC_T_ method and normalized to *ACTIN2* levels. Data are means (±SE) of 2 biological replicates with 1–2 technical replicates. (**C**) GUS activities *in planta* after SA treatment. p*WRKY53*∷GUS lines in Col-0 and *rev5* background, and p*REV*∷GUS plants in *rev9* knock-out mutant [24,33],were used to quantify promoter activities. Data are means (± SE) of 2 biological and 2 technical replicates: 6-13 plants per genotype per condition. Significant differences were analysed using Kruskal–Wallis tests (*: *P* ≤ 0.05; **: *P* ≤ 0.01). See also S4 Table.

While the biosynthesis and catabolism of SA appears to be modulated by *REV* and *WRKY53* (Figs 3A-C), it has also been shown that *WRKY53* expression can be activated by exogenous SA in wild-type plants [22]. To identify whether *REV* expression is also regulated by SA and whether *REV* is involved in SA- induced expression of *WRKY53*, we made quantitative measurements of β-glucuronidase (GUS) activity *in planta* using both *REV* and *WRKY53* GUS-reporter lines after SA treatments. As expected, 19 h after SA-spraying on p*W53*∷GUS rosettes, *WRKY53* expression was clearly induced. These results were modified when p*W53*∷GUS was expressed in the *rev5* background (Fig 3C). Indeed, *WRKY53* expression in this line was weaker and no longer significantly induced by SA when comparing to p*W53*∷GUS line with wild-type REV. This demonstrates that the hormonal regulation of *WRKY53* expression is modulated through the transcriptional factor REV. Moreover, our GUS expression analyses using p*REV*∷GUS plants indicate that *REV* expression itself was not affected by SA treatments (Fig 3C).

### The *REV-WRKY53* interaction controls early and late events of plant development from leaf production to fruit ripening

We performed a comprehensive analysis of plant development from germination to late developmental stages including specific hallmarks such as bolting (flower bud emergence), flowering and mature siliques.

The multivariate effects of the *REV-WRKY53* genetic interaction were first analysed through a principal component analysis (PCA) performed on 14 growth-related traits (S3 Table). First and second principal components (PCs) explained 40.2 % and 32.8 % of the total variance, respectively (S3 Fig). Projection of individuals revealed high genotypic variability as indicated by the distance of the genotypes from the centroid of each other. PC1 was mainly explained by the number and dry mass (DM) of leaves, flowering time and rate of leaf production (RLP; S3 Table), PC2 was driven by senescence- and fitness-related traits, such as the senescent area, reproductive allocation (calculated as the ratio between total silique and rosette DM) and stem size (S3 Table).

Throughout their life cycle, the different mutants exhibited contrasting growth trajectories (Fig 4A). The *rev10-d* gain-of-function and and the *rev5* loss-of-function mutant showed antagonistic growth strategies and presented opposite effects in RLP, total leaf number, flowering time and rosette DM (Figs 4A-D). In addition, *rev10-d* presented a faster and longer production of leaves, with a delayed flowering time compared to Col-0. As a result, *rev10-d* had a higher rosette DM. By contrast, *rev5* displayed a reduced RLP_max_, less leaves, a precocious flowering time and thus a lower rosette DM compared to Col-0 (Figs 4A-D). While *rev10-d* and *rev5* had opposite phenotypes, *wrky53* mutant plants presented similar phenotypes as *rev5* but with a less pronounced effects: although *wrky53* tented to have a higher RLP_max_, it presented a significant lower total leaf number, and less days to reach flowering compared to Col-0. The *wrky53rev5* double mutant displayed similar trends as *rev5* and *wrky53* but with an intermediate phenotype (Figs 4A-D). During fruit ripening, the reproductive allocation was also greatly affected by *REV* and *WRKY53* expression; *wrky53* presented a significantly higher production of siliques per rosette DM than Col-0 whereas *rev10-d*, *rev5* and *wrky53rev5* had lower values (Fig 4E). On the contrary, although *rev10-d* had a much bigger rosette DM, this mutant produced less DM of siliques per rosette DM (Figs 4E-F).

**Fig 4.**
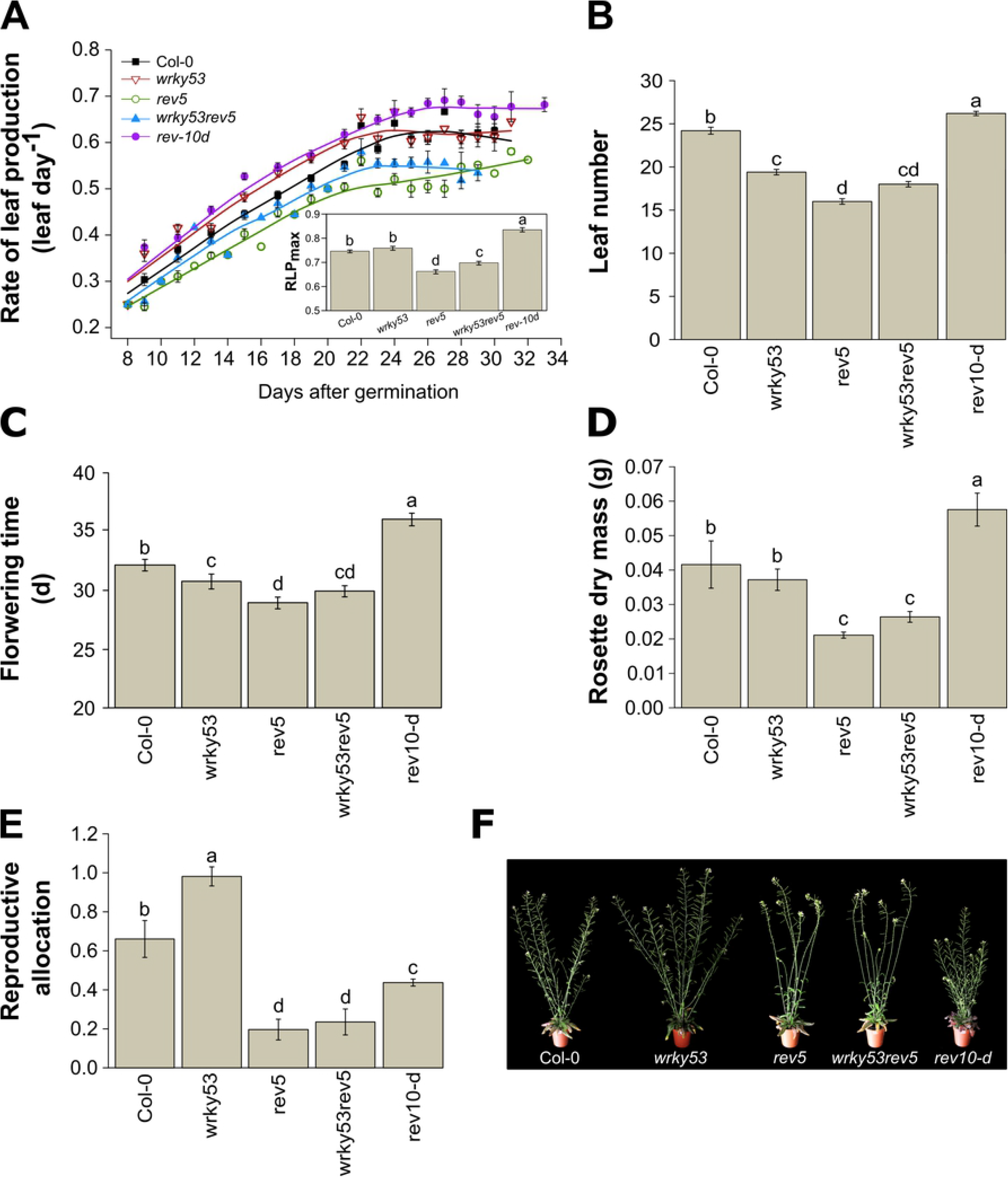
Developmental analysis of the different *REV* and *WRKY53* mutants. **(A)** Rate of leaf production (RLP) during days after germination. Leaf number was counted from the two first true leaves until bolting. Insert in **A** represents the maximum RLP calculated as the slope (leaf per day), estimated by fitting a linear curve for each plant over time. **(B)** Leaf number at flowering. **(C)** Time to reach flowering. **(D)** Rosette dry mass at bolting. **(E)** Reproductive allocation at mature silique stage, calculated by the ratio of total silique dry mass per rosette dry mass. **(F)** A representative picture of rosettes and reproductive organs of the different genotypes at the mature silique. Bolting was defined by the emergence of the flower buds. Flowering was reached when the first flower was open and mature silique stage was considered from the first yellowing siliques. Data are means (± SE) of 5-30 plants per genotype. The different letters indicate significant differences following Kruskal–Wallis tests (*P* ≤ 0.05). See also S2 and S3 Tables.

The analysis of the *REV*-*WRKY53* genetic interaction over the development revealed an interactive effect of the *REV* and *WRKY53* mutations on leaf number, flowering time, rosette DM, and a marginally significant effect on senescence at 10 days after flowering and reproductive allocation (results of two-way ANOVA; S2 Table).

### *REV* and *WRKY53* mediate plant responses to pathogen

Because SA metabolism as well as camalexin production were altered in *wrky53* and *rev* mutants and a general overlap of senescence-associated and pathogen-associated genes has been reported, we aimed to examine pathogen response in the mutants compared to wild-type plants. To explore the crosstalk between the *REV-WRKY53* interaction and plant immunity, we analysed the phenotypic responses to pathogen infection at early developmental stages. For all genotypes, 16-day-old rosettes (without visual senescence symptoms) were infected by spraying a suspension of a genetically modified strain of *Pseudomonas syringae* pv *tomato* DC3000 (*Pst*) constitutively expressing green fluorescent proteins (GFP), that allowed us to monitor its quantification in leaves (Fig 5A). In addition, plant susceptibility was followed after infection using chlorophyll fluorescence (ChlF) imaging through photosynthetic efficiency (*F*_v_/*F*_m_) measurements of whole-rosettes (Fig 5A). Response ratio of *F*_v_/*F*_m_, calculated as the relative ratio of *Pst*-infected plants compared to mock-treated plants, showed that 2 days after infection (DAI) *wrky53* and *rev5* tended to be more sensitive to the infection compared to Col-0, whereas *wrky53rev5* presented a significantly intensified sensitivity (Fig 5B). On the opposite, *rev-10d* appeared to be less sensitive than the others mutants and had a faster recovery of photosynthetic capacities, as illustrated by a reduced response ratio of *F*_v_/*F*_m_ at 6 DAI when comparing to Col-0 (Fig 5B). Investigation of bacterial growth through GFP quantification at 14 DAI indicated that *wrky53rev5* had an increased colonisation of *Pst* in leaves (Fig 5C). These findings were confirmed by independent experiments (S4 Fig) but also through classical *Pst* bacterial leaf infiltration assays with subsequent bacterial growth curve evaluation (Fig 5D). Results of two-way ANOVA at 4 DAI revealed a strong additive effect of *REV* and *WRKY53* mutations (*P* < 0.001 and *P* < 0.01, respectively) on the susceptibility to *Pst* infection (*via* bacterial quantification, Fig 5D, S2 Table).

**Fig 5.**
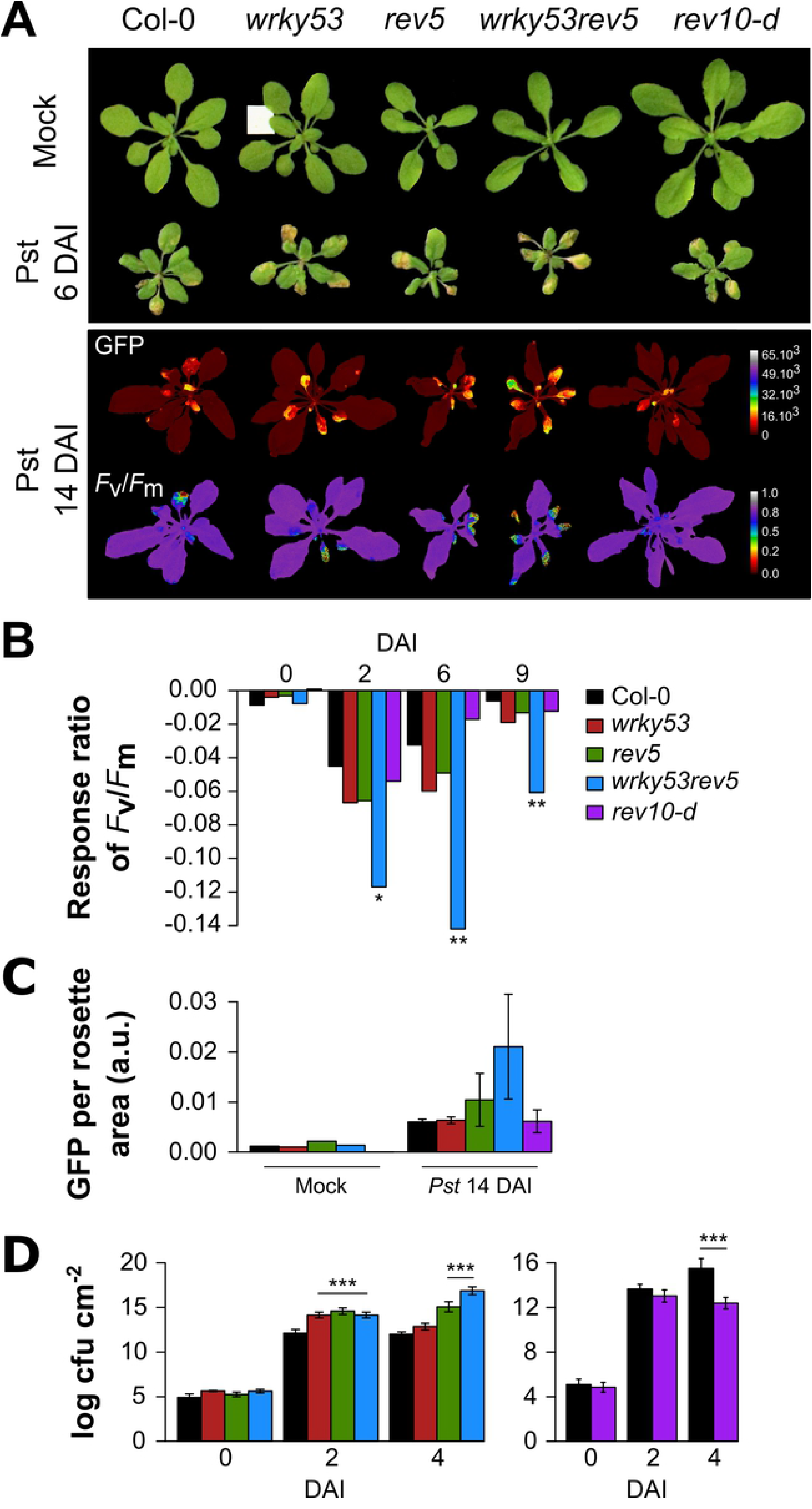
Pathogen assays using *Pseudomonas syringae* inoculation. 16-day-old plants were infected with *Pseudomonas syringae* (*Pst*) pv *tomato* DC3000, constitutively expressing GFP by spraying a *Pst* suspension on leaf surfaces. Bacterial colonisation was followed by quantification of GFP fluorescence *in planta* and plant disease was investigated using chlorophyll fluorescence imaging. **(A) top**: a representative picture of rosettes 6 days after infection (DAI), in parallel to the respective mock condition. **Bottom**: a representation of GFP fluorencence *in planta* and photosynthetic efficiency (*F*_v_/*F*_m_) in *Pst*-infected plants 14 DAI **(B)** Response ratio of *F*_v_/*F*_m_ during DAI, calculated as the relative ratio of *Pst*-infected plants compared to mock-treated plants (*Pst*-Mock)/Mock. Significant differences were analysed using two-way ANOVA (genotype by treatment interaction) on *F*_v_/*F*_m_ values (n = 6-8; *:*P* ≤ 0.05 and **:*P* ≤ 0.001). **(C)** Quantification of bacterial growth *in planta* expressed in GFP units per rosette area compared to the respective mock condition, 14 DAI. Data are means (± SE) of 3 plants. **(D)** Quantification of bacterial growth after infection (cfu = colony forming units). Data are means (± SE) of 4-12 plants. Significant differences were analysed using Kruskal–Wallis tests (***: *P* ≤ 0.001). See also S4 Fig.

### *REV* and *WRKY53* connects plant growth, SA metabolism and immunity

The analyse of correlations amongst traits showed that RLP_max_, flowering time, SA content and *F*_v_/*F*_m_ response ratio after *Pst* infection, were all positively correlated (Fig 6). For instance, RLP_max_ and flowering time were highly correlated (*r*^2^ = 0.94; *P* ≤ 0.05), as well as SA content with *F*_v_/*F*_m_ response ratio (*r*^2^ = 0.95; *P* ≤ 0.05). Moreover, increased RLP_max_ and delayed flowering time explained 86% and 82% of the variation in SA content, respectively. Plant sensitivity to pathogen infection (*i.e. F*_v_/*F*_m_ values) was also explained by developmental traits: 66% by RLP_max_ and 60% by flowering time. *rev10-d* presented an opposite behaviour compared to the knock-out mutants, in which *wrky53rev5* had an intermediate phenotype between *wkry53* and *rev5*.

**Fig 6.**
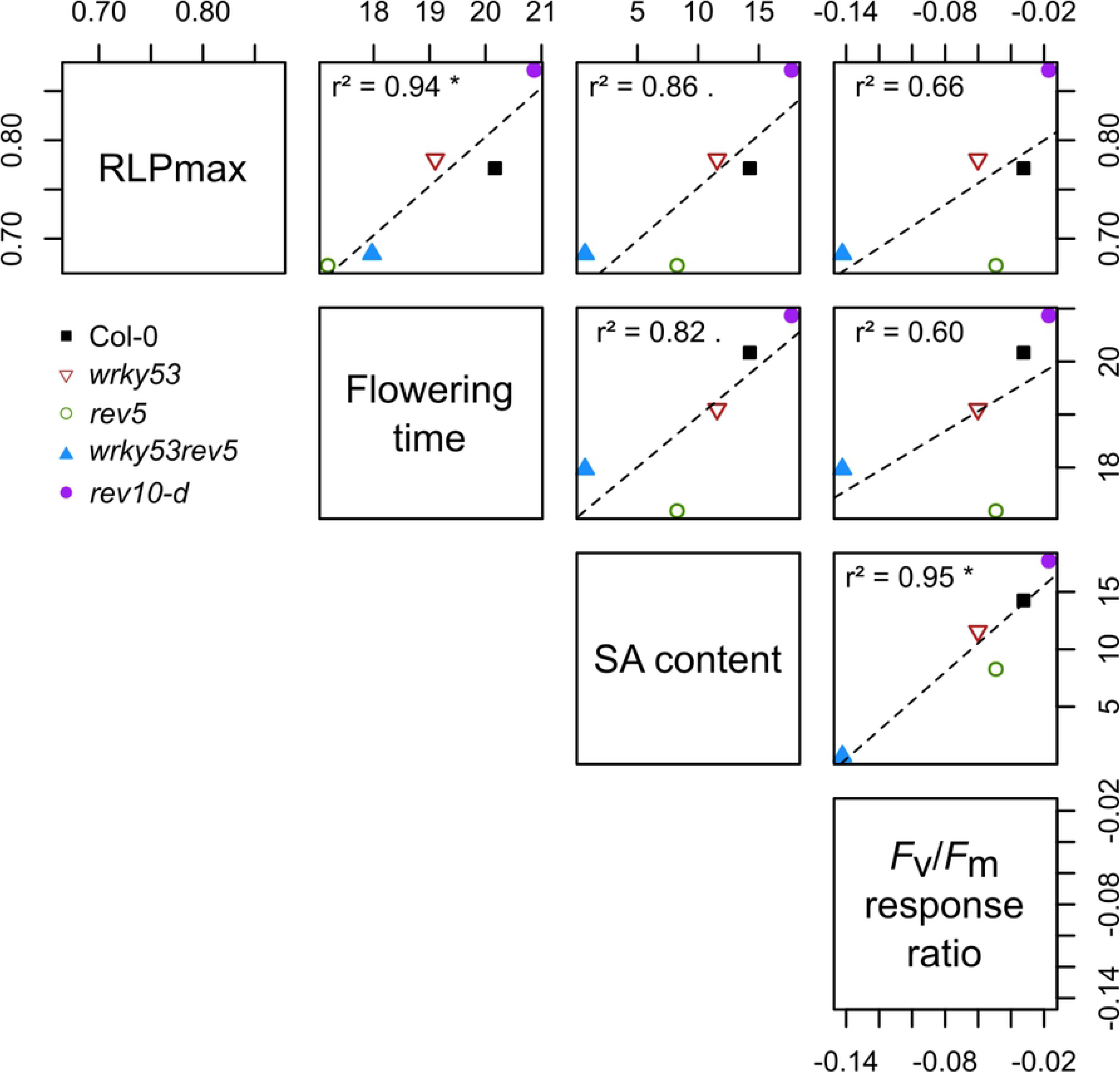
Correlations amongst key traits of plant development and immunity. Pearson correlation coefficients within the different genotypes of key traits of plant development and immunity. RLP_max_ is the maximum rate of leaf production calculated as the slope estimated by fitting a linear curve for each plant over time. Salicylic acid (SA) and *F*_v_/*F*_m_ response ratio were measured at 15 days after flowering and 6 days after inoculation, respectively. (*: *P* ≤ 0.05, .: *P* ≤ 0.01).

## Discussion

The coordination of developmental programs throughout the life cycle of a plant is a central question of plant biology and phenotypic integration. In this study, we showed that the genetic interaction between *WRKY53*, a key senescence-related gene [17], and *REV*, which is known as a leaf patterning gene [35], contribute to the fine-tuning of plant development, immunity and senescence.

The WRKY family represents one of the most abundant groups of TFs involved in the control of senescence [36]. As many TFs, WRKY members are also key actors in integrating developmental processes and stress responses [15]. Consistently, different WRKYs have been shown to be pivotal in pathogen defence [16]. For instance, WRKY53 acts as a positive regulator of basal resistance against *Pseudomonas syringae* (*Pst*) [19,20]. However, the upstream regulation of WRKYs appears to be very complex and remains far from being completely understood. Here, we investigated the pleiotropic role of REV, because previous studies demonstrated that REV directly and positively regulates the expression of *WRKY53* [24]. Moreover, REV also binds to additional differentially expressed senescence-associated genes during leaf senescence [24,30], which suggests that this gene controls many aspects of plant development through its interaction with senescence-related genes. Consistently, we found that the *REV-WRKY53* interaction controls several developmental traits: leaf production rate, flowering time and senescence, which later determine the reproductive success of plants (*i.e*., biomass production and reproductive allocation). Interestingly, multivariate analysis of developmental and senescence traits revealed that traits related to flowering time and vegetative growth were strongly correlated, shaping the first dimension of phenotypic variation (*e.g*., PC1 explained more than 40% of total phenotypic variance). This is consistent with previous analyses of natural variability performed in *A. thaliana*, in which traits related to growth rate and life cycle duration also represents the first axis of phenotypic variability [37]. By contrast, we found that SAGs were correlated to fitness-related traits, such as stem length and DM, as well as reproductive allocation. This suggests that senescence is a key process involved in the control and efficiency of reproduction, and that its role on fecundity dominates the role of flowering time or vegetative growth. *REV* mutants exhibited stronger effect on senescence and fitness traits than *WRKY53* mutants, which were more variable along the first phenotypic dimension. Strikingly, this finding reveals that *REV*, a gene known as a master regulator of early leaf patterning [26–28], critically controls late developmental phases and reproduction while, inversely, *WRKY53*, a gene known as a master regulator of late leaf development [38], determines vegetative growth even at early developmental stages. This unanticipated result demonstrates how late and early developmental events are tightly intertwined by molecular hubs that interact with each other.

Moreover, our study revealed that the plant’s metabolism during growth was modified by *REV* and *WRKY53*. Amongst the studied metabolites, SA and its related components were the most variable ones between the mutants and Col-0. The analysis of the interaction between *REV* and *WRKY53* by two-way ANOVA indicated that both genes were independently involved in SA-related metabolic changes, and with a stronger effect of *REV*. By contrast, *REV* and *WRKY53* exhibited significant interaction in the control of plant development, notably for total leaf number at reproduction, flowering time and shoot dry mass. In addition, *REV* and *WRKY53* exhibited no or marginal interaction in the control of senescence and fitness traits. Taken together, our results suggest that REV and WRKY53 strongly interact during plant development, while they later act independently in the control of senescence through the SA signalling pathway.

Since we found that the biosynthesis as well as catabolism of SA can be modulated by *REV* and *WRKY53*, SA-related gene expression was analysed. Two SA biosynthetic pathways have been characterized in plants, the phenylalanine ammonia lyase (PAL) pathway and the isochorismate (IC) pathway, both using chorismite as primary metabolite. However, approx. 90% of the SA produced after pathogen attack or UV light exposure are produced via the IC pathway [39,40]. Isochorismate synthase (ICS) and isochorismate pyruvate lyase can convert chorismate to SA via isochorismate. Moreover, SA can also be stored in form of inactive SA glycosides (SA 2-O-β-D-glucose and SA glucose ester) which are actively transported to the vacuole and can be converted back to SA [41,42]. 2,3- and 2,5-dihydroxybenzoic acid (2,3-DHBA and 2,5-DHBA) sugar conjugates appear to be the major storage form in vacuoles of old Arabidopsis leaves [32,43] and SA 3-hydroxylase (S3H), which converts SA to 2,3-DHBA, prevents over-accumulation of active SA implying an important role in regulating leaf senescence [32]. Consistent with previous findings [42], we showed that the expression of the *S3H* gene increased in senescing leaves of wild-type Col-0 plants. However, this up-regulation was damped in the mutants, with a significant interaction of *REV* and *WRKY53* mutations. In addition, we found that *ICS1* expression is also diminished in the mutants compared to the senescence-associated increase in wild-type, with a strong decrease in *rev10-d*. Thus, the control of isochorismate synthase and SA 3-hydroxylase activity by the *REV-WRKY53* interaction is expected to be crucial for the fine-tuning of plant senescence.

However, SA is known as a multifaceted hormone which is also of utmost importance for disease resistance, but also for flowering and senescence [44,45]. Through its effect on the SA signalling pathway, the *REV*-*WRKY53* interaction could be a key driver of the already described growth-defence trade-off [10]. In order to test this hypothesis, we sprayed plant seedlings with *Pst* to test the response of *REV* and *WRKY53* mutants to pathogen attack. Since *Pst* enters the plant through natural openings or wounds, this method just mimicked their natural entry into the apoplastic space [46]. Interestingly, we observed improved resistance against these pathogens in *rev1*0*-d* compared to Col-0, although this mutant grew faster and higher than the wild-type. This suggests that the behaviour and phenotype of *rev10-d* can be explained by a higher SA content but also by a faster and longer production (*i.e*., delayed flowering) of healthy leaves after infection. In addition, not only SA metabolism but also the production of camalexin, which appears to be the major phytoalexin involved in biotic responses in *A. thaliana* [31,46], is also altered in the mutants. Camalexin production exhibited the same pattern as free SA, most likely contributing to the resistance of *rev10-d*.

Moreover, we have shown that the induction of *WRKY53* expression by SA is dependent on REV as SA induction of a *WRKY53* promoter driven GUS reporter gene was severely impaired in the *rev* mutant background. However, *REV* expression appeared to be insensitive to SA, which means that there is a feed-back loop of SA production to *WRKY53* expression but not to REV expression. In the same line of evidence, REV is also involved in the response of *WRKY53* expression by H_2_O_2_ [24] and SA induces the accumulation of H_2_O_2_, and *vice versa*. On the other hand, the DNA-binding activity of the REV protein is redox-sensitive indicating a very complex feed-back regulation between *REV*, *WRKY53*, H_2_O_2_ and SA.

Collectively, our results indicate that the genetic interaction between *REV* and *WRKY53* seems to be strongly dependent on the developmental stage and that *REV* acts upstream of *WRKY53* to modulate SA signalling from early to late plant development.

## Materials and Methods

### Plant material and growth conditions

The *A. thaliana* ecotype Col-0 and the following mutants in the Col-0 background were used in this study: *rev5*, (A260V) a strong ethyl-methylsulfonate (EMS) allele of *REV*; *wrky53*, (SALK-034157) a T-DNA insertion line in the second exon of *WRKY53* [22]; *wrky53rev5*, a double knock-out mutant [24] and *rev10-d*, a semi-dominant gain-of-function of *REV* allele where *REV* mRNA was rendered resistant to the negative regulation by microRNAs [27]. p*WRKY53*∷GUS lines in Col-0 and *rev5* background, and p*REV*∷GUS plants in *rev9* knock-out mutant [24,35] were used to quantify promoter activity *in planta*. Plants were grown on standard soil (9:1 soil and sand) under controlled conditions: in long days (16 h day; 8 h night), low light (~70-80 μE m^−2^ s^−1^ at plant height) and an ambient temperature of 21 °C (see [47] for details). Three independent experiments were done for developmental analysis, including senescence and plant productivity quantification.

### Plant developmental analysis

The number of leaves that were visible to the naked eye (at least/minimum of 2-3 mm) was counted every day to determine the RLP from the emergence of the first two true leaves until macroscopic visualization of flower buds (bolting). The maximum RLP was determined as the slope estimated by fitting a linear curve for each plant over time. Bolting, flowering time and mature silique were determined as the number of days from germination until bolting, the first flower open and the first yellowing siliques, respectively. At each stage, plants were individually harvested and leaf blades were separated from their petiole considering the position (age) within the rosette. Leaf blades were scanned for measurements of area using the ImageJ software (1.47v, Rasband, Bethesda, Maryland, USA). The maximum rate of leaf expansion (R_max_) was determined as for [49]. Specific leaf area (SLA) was calculated as the ratio of total leaf area to leaf DM. DM of the different organs (leaves, petioles and stems) was measured by placing them in separate paper bags at 67 °C for at least 5 days.

### Senescence and plant productivity quantification

Due to significant differences in phenology of mutants, senescence progression was analysed from the first flower open for each individual plant. Senescence was quantified in leaves using ChlF imaging (Imaging-PAM; Maxi version; ver. 2-46i, Heinz Walz GmbH; see [47,48]). The maximum quantum yield of photosystem II (PSII) was estimated by the ratio of variable to maximal ChlF (*F*_v_/*F*_m_, also called photosynthetic efficiency) on dark-adapted plants, after 15-20 min. Senescent area was estimated by a clustering approach of *F*_v_/*F*_m_ values of each pixel of the ChlF image (according to [49]). Senescent area was calculated by the ratio between pixel number for *F*_v_/*F*_m_ ≤ 0.6 and the total pixel number of the rosette (or in single leaves). Silique number was counted at mature silique stage and siliques were dried in a paper bag at 67 °C for at least 5 days. Reproductive allocation was calculated by the ratio of total silique DM per rosette DM.

### Metabolomics analysis: extraction of primary and secondary metabolites

For each genotype and senescence stage 5 replica samples were created. For each replica, leaves at position 5 to 11 were po-oled and weighed for each plant from flowering to mature silique stages. 10 mg of freeze dried, retched (5 mm steel ball; 30 sec) plant material was extracted with 200 μl 80 % methanol (MeOH; 0.1 % formic acid (FA); ice cold). The pellet was re-extracted with 200 μl 20 % MeOH (0.1 % FA; icecold) and both supernatants were combined. The whole extraction process was done at 10 °C including a 10 min sonication and centrifugation step (18600 g). For the analysis of the free and conjugated phytohormones the final extract was directly used for targeted liquid chromatography–mass spectrometry (LC-MS) analysis (see Supplemental methods). To determine the amount of bound SA and DHBA, 50 μl of the plant extracts were hydrolyzed with 4 μl concentrated FA (99 °C; 2 h; 800 rpm), cooled down on ice and centrifuged (10 °C; 18600 g) before submitting them into the targeted LC-MS pipeline. For the analysis of sugars and amino acids, 100 μl extracts were dried down in a speed vacum concentrator before derivatisation for gas chromatography-MS (GC-MS) analysis (see Supplemental methods). For non-targeted LC-MS analysis, the same extraction was performed as for the targeted approach (see Supplemental methods). The combined supernatants from the extraction were dried down in a speed vacum concentrator and afterwards redissolved in 100 μl 20 % MeOH containing 0,1 % FA and 9 μM L-Encephalin as an internal standard. In addition 10 μl of each sample was combined to create a pool sample of the entire analysis. 5 μl of any sample were injected for the non-targeted LC-MS analysis.

### Gene expression analysis

In two independent experiments (same growth conditions as above), gene expression levels were followed by qRT-PCR according to [48]. Leaves at position 10 were harvested from flowering to mature silique stages in individual plants. A pool of 5 leaves per genotype/stage was used for RNA extraction and cDNA synthesis. Samples harvested at mature silique stage were removed from the analyses because very low amounts of RNA could be isolated and no reliable transcript levels were detected. Transcriptional changes were calculated based on the comparative ΔΔC_T_ method [50] and normalized to *ACTIN2* levels. Primers used are listed in S4 Table.

### Quantitative measurement of GUS activity in *planta* after SA treatments

Two-week-old plants were used to analyse GUS activity *in planta* by spraying whole-rosette with 2 mM SA in 0.15 % ethanol and 0.02 % Silwet L-77. Control plants were treated using appropriate mock solution. GUS activity was measured according to [51]. Rosettes were individually frozen in liquid nitrogen, homogenized using Mill MM 200 (Retsch GmbH, Germany) and resuspended in 1 mL GUS Extraction Buffer (50 mM sodium phosphate pH 7.0, 10 mM EDTA, 10 mM β-mercaptoethanol, 0.1 % triton X-100, 0.1 % sarcosyl and 25 μg ml^−1^ PMSF). Samples were strongly vortexed and centrifuged at 13.000 rpm for 15 min at room temperature. 200 μL of supernatant were then kept on ice. 10 μL of extracts were added in 130 μL Assay Buffer (GUS Extraction Buffer containing 2 mM 4-methylumbelliferyl β-D-glucuronide; MUG). After 20 min at 37 °C, 10 μL of the reaction were transferred to 190 μL of a 200 mM sodium carbonate solution in a 96-well dark plate. Fluorescence was measured on a TriStar LB941 plate reader (Berthold Technologies, Germany) at 460 nm (emission) and 355 nm (excitation). A standard curve from 0 to 50 μM of 4-methylumbelliferone (4-MU) was used to calculate moles of MU min^−1^ produced after the cleavage of MUG by GUS enzyme in *planta*. 2 μL of extracts in 200 μL of 5-fold diluted Bradford Roti-Quant (Roth, Germany) were used to quantify protein concentration of samples [52] and to calculate the final GUS activity in nmol MU min^−1^ mg^−1^.

### Pathogen assays and bacterial quantification

Pathogen assays were performed using *Pst* pv *tomato* DC3000 *att*Tn*7*-egfpmut3 constitutively expressing GFP. *Pst* pv *tomato* DC3000 was labeled via Tn*7* site-specific transposition with the mini-Tn*7* vector pURR25 [53] essentially as described in [54]. Bacteria were grown overnight at 28 °C in dark in specific medium (1 % (w/v) tryptone, 0.6 % (w/v) yeast extract, 0.15 % (w/v) K_2_HPO_4_, 0.06 % (w/v) NaCl, pH 7) supplemented with rifampicin (50 μg mL^−1^) and kanamycin (25 μg mL^−1^) to OD_600_ = 0.8. The inoculum preparation and Arabidopsis infection were then done as described by Katagiri et al., (2002). A final density of 5×10^8^ colony-forming unit mL^−1^ (cfu mL^−1^) of bacterial suspension, in sterile water with 0.02 % (v/v) Silwet L-77, was used for spray inoculation of 16-day-old plants. As control, a mock solution (0.02 % (v/v) Silwet L-77 in sterile water) was used. Inoculation was done twice (8h intervals) and plants were kept covered with a transparent plastic lid for 2 days. Bacterial colonisation was analysed by quantifying GFP fluorescence in abaxial leaf surface using a Typhoon FLA9500 laser scanner (GE Healthcare) and ImageJ analysis. Plant disease was followed after the infection using ChlF imaging through *F*_v_/*F*_m_ measurements of whole-rosettes, as described above. A response ratio of *F*_v_/*F*_m_ was calculated as the relative ratio of *Pst*-infected plants compared to mock-treated plants followed (*Pst*-Mock)/Mock. In independent experiments, *Pst* pv *tomato* DC3000, grown overnight at 28 °C to OD_600_ = 0.2 in King’s B medium supplemented with 50μg mL^−1^ of rifampicin, was infiltrated at a density of 1×10^4^ cfu mL^−1^ in 10 mM MgCl_2_ in leaf tissue with a needleless syringe. Middle age leaves (fourth or fifth leaves from young to old) of 6-week-old plants, grown under short day conditions, were used for infiltration. Bacterial growth (cfu cm^2^) was measured in two discs per leaf after extracting bacteria. For this purpose, leaf discs were incubated in 70 % ethanol for 1 min, dried on filter paper, subsequently washed briefly washing in water for 1 min, again dried on filter paper and then homogenized in 200 μL of 10 mM MgCl_2_. Subsequent serial dilutions and depositions on LB-plates with 50 μg mL^−1^ of cycloheximide and 50 μg mL^−1^ of rifampicin were performed by using multichannel pipettes.

### Statistical analyses

PCAs were performed on developmental or metabolite values to focus on genotypic effects at a given developmental stage [55]. *REV* and *WRKY53* mutation effects were analysed in analyses of variance (ANOVA) on PC coordinates. Metabolite patterns (heat map) of mutants were analysed using response ratio, calculated between log-transformed values of Col-0 and a mutant at a given stage. A hierarchal cluster diagram was constructed using Manhattan distances and Ward’s hierarchical clustering method. Comparisons of mean trait values between genotypes were performed using Kruskal–Wallis nonparametric tests for developmental analysis. All analyses were performed using R software (3.4.3 v.; R Development Core Team, 2012).

### Accession numbers

Sequence data from this article can be found in the GenBank/EMBL data libraries under the following accession numbers: *ACTIN2* (At3g18780), *ICS1* (*SID2*; At1g74710), *REV* (At5g60690), *S3H* (*SAG108*; At4g10500), *WRKY53* (At4g23810).

## Acknowledgment

We thank Manuela Freund, Julian Fratte and Gesine Seibold for their technical support. We thank Brian H. Kvitko and Emanuele Scacchi for providing us the line of *Pseudomonas syringae* pv *tomato* DC3000 attTn7-egfpmut3 constitutively expressing GFP.

## Author Contributions

J.B. designed the research, performed experiments, analyzed the data, wrote the manuscript and secured funding. U.Z. designed, supervised the research, wrote the manuscript and secured funding. S.W. designed, supervised the research, contributed to writing and secured funding. J.D. performed qRT-PCRs for gene expression analysis, and contributed to writing. F.V. wrote the manuscript. M.S., E.v.R.L., J.K. and B.S. performed metabolite quantification. J.M.K. designed and provided the line of *Pseudomonas syringae* pv *tomato* DC3000 *att*Tn*7*-egfpmut3 constitutively expressing GFP. D.K. performed bacterial leaf infiltration assays. All co-authors read and approved the final manuscript. JB was supported by the Institutional Strategy of the University of Tuebingen (Deutsche Forschungsgemeinschaft, ZUK 63) and also funded by the Alexander von Humboldt Foundation. This work was supported by the Deutsche Forschungsgemeinschaft (CRC1101, B06).

## Supporting information

**S1 Table.**
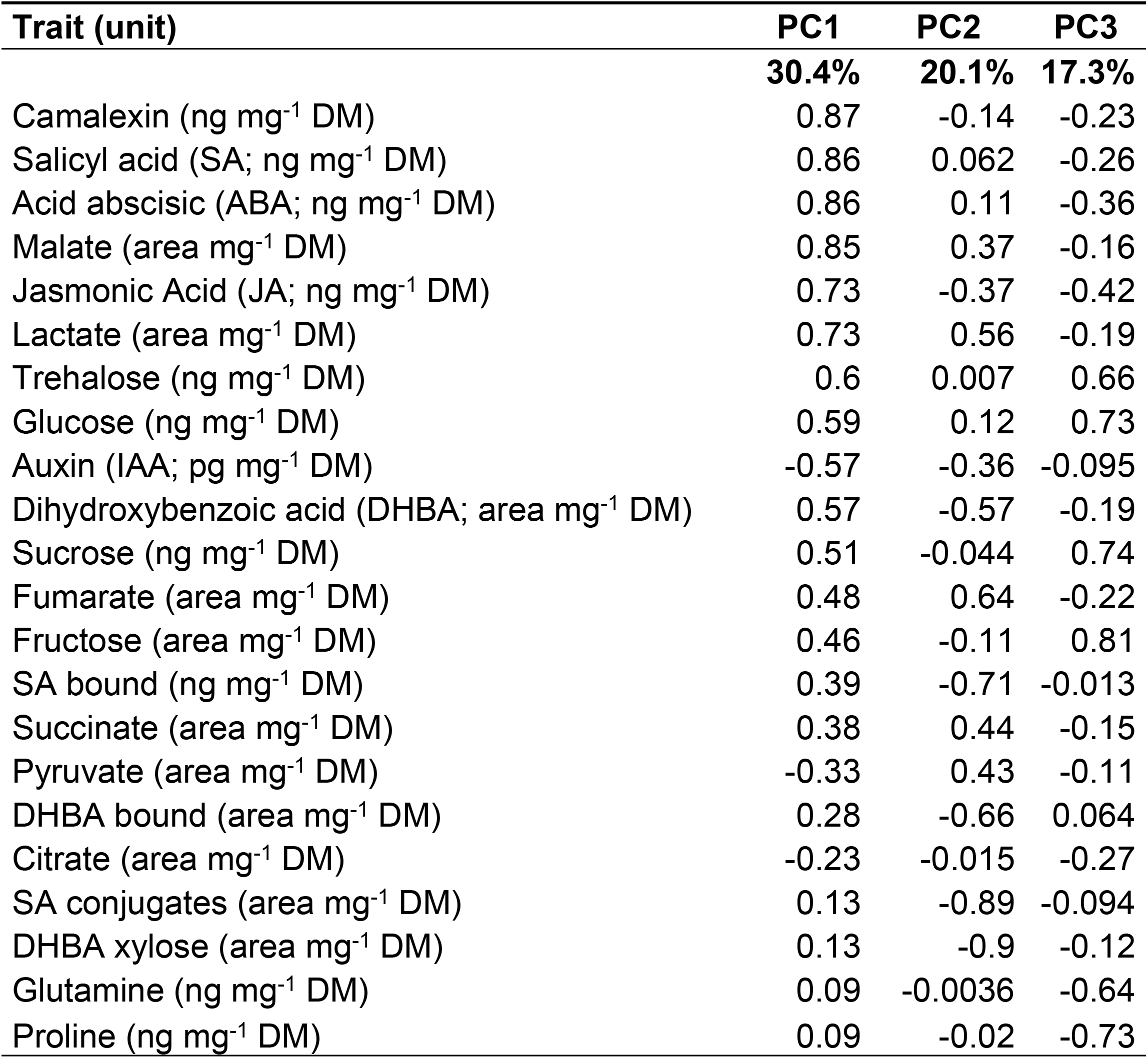
Loadings of the variables included in the PCA on mean of metabolite abundance values. Percentages indicate the percentage the total variance explained in the three first principal components (PC). Loadings are correlation coefficients between the variables and PCs. DM = dry mass

| Trait (unit) | PC1 | PC2 | PC3 |
| --- | --- | --- | --- |
|  | **30.4%** | **20.1%** | **17.3%** |
| Camalexin (ng mg<sup>-1</sup> DM) | 0.87 | -0.14 | -0.23 |
| Salicyl acid (SA; ng mg<sup>-1</sup> DM) | 0.86 | 0.062 | -0.26 |
| Acid abscisic (ABA; ng mg<sup>-1</sup> DM) | 0.86 | 0.11 | -0.36 |
| Malate (area mg<sup>-1</sup> DM) | 0.85 | 0.37 | -0.16 |
| Jasmonic Acid (JA; ng mg<sup>-1</sup> DM) | 0.73 | -0.37 | -0.42 |
| Lactate (area mg<sup>-1</sup> DM) | 0.73 | 0.56 | -0.19 |
| Trehalose (ng mg<sup>-1</sup> DM) | 0.6 | 0.007 | 0.66 |
| Glucose (ng mg<sup>-1</sup> DM) | 0.59 | 0.12 | 0.73 |
| Auxin (IAA; pg mg<sup>-1</sup> DM) | -0.57 | -0.36 | -0.095 |
| Dihydroxybenzoic acid (DHBA; area mg<sup>-1</sup> DM) | 0.57 | -0.57 | -0.19 |
| Sucrose (ng mg<sup>-1</sup> DM) | 0.51 | -0.044 | 0.74 |
| Fumarate (area mg<sup>-1</sup> DM) | 0.48 | 0.64 | -0.22 |
| Fructose (area mg<sup>-1</sup> DM) | 0.46 | -0.11 | 0.81 |
| SA bound (ng mg<sup>-1</sup> DM) | 0.39 | -0.71 | -0.013 |
| Succinate (area mg<sup>-1</sup> DM) | 0.38 | 0.44 | -0.15 |
| Pyruvate (area mg<sup>-1</sup> DM) | -0.33 | 0.43 | -0.11 |
| DHBA bound (area mg<sup>-1</sup> DM) | 0.28 | -0.66 | 0.064 |
| Citrate (area mg<sup>-1</sup> DM) | -0.23 | -0.015 | -0.27 |
| SA conjugates (area mg<sup>-1</sup> DM) | 0.13 | -0.89 | -0.094 |
| DHBA xylose (area mg<sup>-1</sup> DM) | 0.13 | -0.9 | -0.12 |
| Glutamine (ng mg<sup>-1</sup> DM) | 0.09 | -0.0036 | -0.64 |
| Proline (ng mg<sup>-1</sup> DM) | 0.09 | -0.02 | -0.73 |

**S2 Table.** Coefficients of genetic interaction between *REV* and *WRKY53* on metabolic, developmental and immune traits. Two-way ANOVA performed on log_10_-transformed values of developmental traits over the developement. ns = non significant, .: P ≤ 0.1, *: *P* ≤ 0.05, **: *P* ≤ 0.001, and ***: *P* ≤ 0.0001. B = bolting, F = flowering, F+10d = 10 days after flowering and S = mature silique stage; SA bound = total SA after hydrolysis - free SA, SA conjugates = SA-(C6)-glycosides, DHBA = dihydroxybenzoic acid; DHBA bound = Total DHBA after hydrolysis - free DHBA; cfu = colony forming units

| Trait | Intercept | REV*WRKY53 |  |
| --- | --- | --- | --- |
| *Developmental analysis* |  |  |  |
| Rate of leaf production | -0.157 *** | 0.013 | ns |
| Leaf number | 1.140 *** | 0.055 | ** |
| Flowering time | 1.555 *** | 0.029 | * |
| Senescent area (F+10d) | 1.158 *** | 0.482 | . |
| Shoot dry mass (S) | -0.854 *** | 0.139 | * |
| Reproductive allocation | -0.358 *** | -0.206 | . |
| Number Silique | 1.883 *** | -0.015 | ns |
| *Metabolic analysis* |  |  |  |
| SA | 8.84 *** | 3.28 | ns |
| SA bound | 22.5 *** | 4.82 | ns |
| SA conjugates | 75556 ** | -5610 | ns |
| DHBA xylose | 3902.6 ** | 140.4 | ns |
| DHBA bound | 265439 *** | 91354 | ns |
| *Pathogen assays* |  |  |  |
| log cfu cm-2 | 16.86 *** | 0.91 | ns |

**S3 Table.** Loadings of the variables included in the PCA on mean of 14 growth-related traits values. Percentages indicate the percentage the total variance explained in the three first principal components (PC). Loadings are correlation coefficients between the variables and PCs.

| Trait (unit) | PC1 | PC2 | PC3 |
| --- | --- | --- | --- |
|  | **40.2%** | **32.8%** | **9.5%** |
| Leaf number (leaf) | -0.94 | 0.05 | 0.02 |
| Leaf dry mass (g) | -0.94 | 0.08 | 0.25 |
| Bolting time (d) | -0.86 | -0.0039 | 0.14 |
| Flowering time (d) | -0.85 | -0.38 | 0.0062 |
| Specific leaf area (SLA; m<sup>2</sup> g<sup>-1</sup>) | 0.78 | 0.46 | -0.23 |
| Whole-rosette area (cm<sup>2</sup>) | -0.76 | 0.5 | 0.14 |
| Rate of leaf production (RLP; leaf d<sup>-1</sup>) | -0.74 | 0.04 | -0.063 |
| Maximum rate of leaf expansion (R<sub>max</sub>; cm<sup>2</sup> d<sup>-1</sup>) | 0.52 | 0.31 | 0.38 |
| Senescent area (%) | -0.27 | 0.62 | -0.68 |
| Stem dry mass (g) | -0.23 | 0.85 | 0.29 |
| Petiole dry mass (g) | -0.19 | -0.85 | -0.19 |
| Stem length (cm) | -0.11 | 0.91 | 0.17 |
| Reproductive allocation | 0.26 | 0.9 | 0.18 |
| Whole-rosette $F_v/F_m$ | 0.37 | -0.64 | 0.62 |

**S4 Table.**
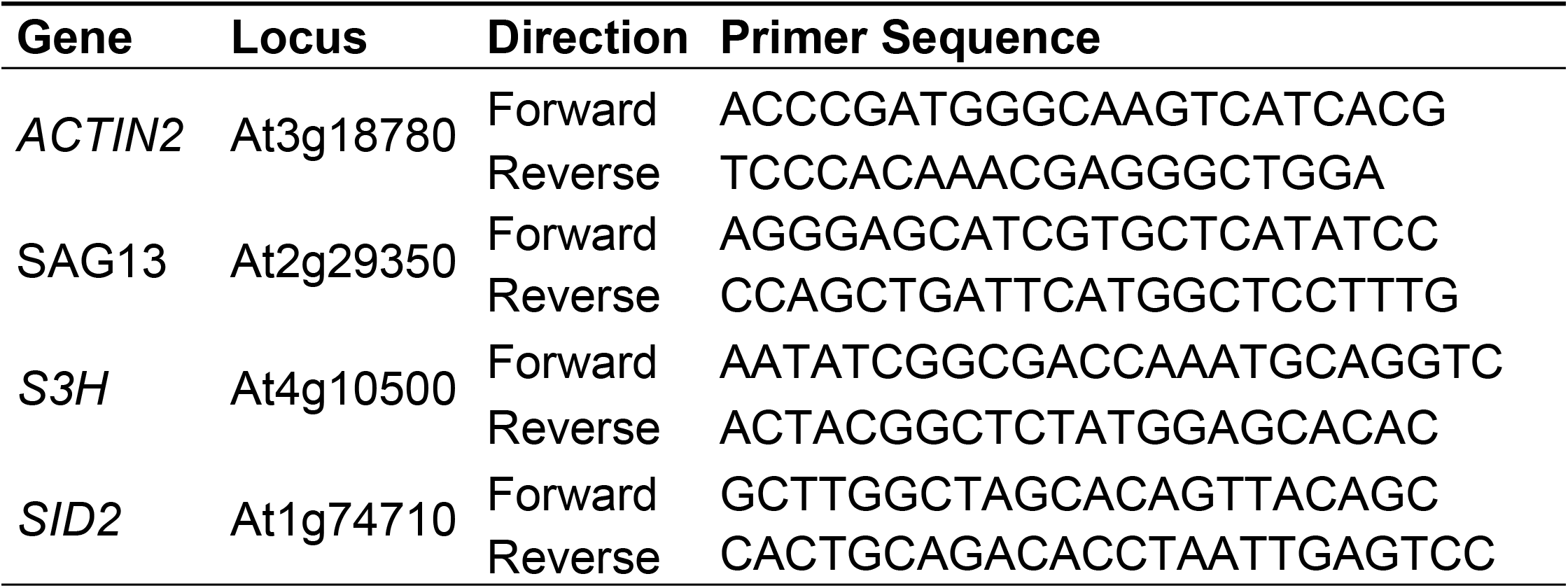
List of primers used.

**S1 Fig. Percentage of senescent area by leaf position of the different mutants compared to Col-0**. (**A**) Percentage of senescent area displayed at 10 days and (**B**) 15 days after flowering. Senescent area was calculated by the ratio between pixel number for *F*_v_/*F*_m_ ≤ 0.6 and the total pixel number of the rosette. Black line means wild-type Col-0, colours show mutants. Data are means (± SE) of 5 plants.

**S2 Fig. Abundances of main metabolites explaining the most significant effects in PCA at 15 days after flowering**. (**A**) camalexin, (**B**) salicylic acid (SA), (**C**) abscisic acid (ABA), (**D**) malate, (**E**) jasmonic acid (JA), (**F**) dihydroxybenzoic acid (DHBA) xylose and (**G**) SA conjugates. Data are means (±SE) of 5 plants. Different letters indicate significant differences between means following Kruskal-Wallis tests (*P* < 0.05). DM = dry mass; SA conjugates = SA-(C6)-glycosides.

**S3 Fig. Principal component analysis on multiple growth-related traits measured on** *REV* **and***WRKY53* **mutants. (A**) Representation of the variables, measured at 10 days after flowering, on the two first principal components. DM = dry mass; RLP = rate of leaf production; R_max_ = maximum rate of leaf expansion; SLA = specific leaf area; *F*_v_/*F*_m_ = maximum quantum yield of photosystem II. (**B**) Projection of individual plants with centres of gravity per genotype (n = 5). Ellipses represent inertia ellipses, centred on the means for each genotype. Their width and height are given by 1.5 times the standard deviation of the coordinates on axes, and the covariance sets the slope of the main axis [58]. *rev5* and *wrky53* are single knock-out of the *REV* and *WRKY53* genes, respectively. *wrky53rev5* is double knock-out (Xie et al., 2014). *rev10-d* a semi-dominant gain-of-function of *REV* allele where *REV* mRNA is rendered resistant to the negative regulation by microRNAs [27].

**S4 Fig. Pathogen assays**. 16-day-old plants were infected with *Pseudomonas syringae* (*Pst*) pv *tomato* DC3000, constitutively expressing GFP by spraying a *Pst* suspension on leaf surfaces. (**A**) Response ratio of photosynthetic efficiency (*F*_v_/*F*_m_) during days after infection (DAI), calculated as the relative ratio of *Pst*-infected plants compared to mock-treated plants (*Pst*-Mock)/Mock. **(B)** Quantification of bacterial growth *in planta* expressed in GFP units per rosette area compared to the respective mock condition. Data are means (± SE) of 3-5 plants per condition.

